# Medical-Grade Mānuka Honey and Mānuka Honey Extract Inhibit Mast Cell Degranulation through inhibition of MRGPRX2 expression: Potential Intravesical Agent for the Management of Interstitial Cystitis/Bladder Pain Syndrome?

**DOI:** 10.64898/2026.08.02.742332

**Authors:** Omar K. A. Abdelwahab, Kamaluddeen Garba, Laurie Lau, David A. Johnston, Andrew F. Walls, Hannah Markham, Brian R. Birch, Jackie C. Evans, Troy L. Merry, Bashir A. Lwaleed

## Abstract

**Rationale:** Neurogenic inflammation is recognised as an important contributor to the pathophysiology of Interstitial Cystitis/Bladder Pain Syndrome (IC/BPS). Substance P (Sub P), a neuropeptide released from sensory nerves, is a potent inducer of mast cell degranulation through the Mas-related G protein-coupled receptor member X2 (MRGPRX2), resulting in the release of pro-inflammatory mediators that perpetuate chronic bladder inflammation. Medihoney, a medical-grade Mānuka honey, possesses well-established antimicrobial and anti-inflammatory properties, and we have recently demonstrated its ability to stabilise mast cells through inhibition of histamine release. However, its effects on Sub P-induced mast cell activation and MRGPRX2-mediated neurogenic inflammation have not previously been investigated.

**Aim of the study:** We aimed to investigate the inhibitory effects of Medihoney and a sugar-free Mānuka honey extract on Substance P-induced mast cell degranulation and MRGPRX2 activation as potential therapeutic approaches for chronic neurogenic inflammation associated with IC/BPS. In addition, we examined the expression of MRGPRX2 in bladder biopsies from patients with IC/BPS.

**Materials and methods:** Human LAD2 mast cells were stimulated with Substance P (1 μM) for 40 minutes following 20-minute pre-incubation with Medihoney or a sugar-free Mānuka honey extract. Mast cell degranulation was quantified by measuring β-hexosaminidase release. MRGPRX2 activation was assessed by intracellular calcium imaging using Fluo-4 in MRGPRX2-expressing HEK-293 cells. Bladder biopsies obtained from patients with IC/BPS and healthy controls were immunostained for mast cell tryptase, chymase and MRGPRX2.

**Results:** Medihoney at 2% and 4% markedly inhibited Substance P-induced mast cell degranulation in LAD2 cells by approximately 90%, an effect that was similarly observed with the sugar-free Mānuka honey extract. Both preparations produced a dose-dependent inhibition of Substance P-induced intracellular signalling in MRGPRX2-expressing HEK-293 cells, demonstrating suppression of MRGPRX2 activation. Furthermore, immunohistochemical analysis of bladder biopsies revealed that approximately 66% of tryptase-positive mast cells expressed MRGPRX2 in patients with IC/BPS, which was significantly higher than that observed in healthy control tissues (25%).

**Conclusion:** The present study demonstrates that mast cells within IC/BPS bladder tissue express increased levels of MRGPRX2, suggesting enhanced responsiveness to Substance P and supporting a role for neurogenic inflammation in the pathophysiology of IC/BPS. Medihoney and the sugar-free Mānuka honey extract significantly inhibit Substance P-induced mast cell degranulation through modulation of MRGPRX2-mediated intracellular signalling, highlighting their potential as novel therapeutic agents for reducing neurogenic bladder inflammation associated with IC/BPS.

**Impact:** This study provides evidence that MRGPRX2-mediated neurogenic mast cell activation is enhanced in IC/BPS and demonstrates, for the first time, that Medihoney and a sugar-free Mānuka honey extract effectively inhibit Substance P-induced mast cell degranulation through modulation of MRGPRX2 signalling. These findings provide new mechanistic insight into the anti-inflammatory actions of Mānuka honey-derived preparations and identify MRGPRX2 as a potential therapeutic target in IC/BPS. The observed inhibition of neurogenic mast cell activation suggests that these naturally derived preparations may offer a novel strategy for limiting chronic bladder inflammation. Overall, this work provides a foundation for future preclinical and clinical studies evaluating the safety and therapeutic efficacy of Medihoney and Mānuka honey-derived compounds in patients with IC/BPS.

## Introduction

Interstitial cystitis/bladder pain syndrome (IC/BPS) is a chronic debilitating inflammatory condition associated with chronic bladder pain and urge to void as well as increased day-and night-time urination frequency (1). Recent reports show a progressive increase in the prevalence of IC/BPS (i.e. 52 to 500/100,000 in females and 8 to 41/100,000 in males), accompanied by an increase in the incidence rate across genders (2, 3). The incidence is estimated at 1.2/100,000. A recent report from the UK highlighted the significant burden that IC/BPS places on patients’ quality of life, work productivity and how it drains healthcare reserves (4). These include a significantly impaired Health-Related Quality of Life (HRQOL) score, greater work productivity loss (41.7%), and, crucially, a higher number of IC/BPS patients attending GP surgeries, outpatient clinics and A&E Departments together with increased inpatient admissions (5). The above statistics are reflected in the finding that the number of patients with IC/BPS who commit suicide is four times that of healthy individuals (6).

The management of IC/BPS presents a notable burden on health services in the UK and worldwide. For example, in the USA, it has been calculated that the mean annual costs associated with the management of IC/BPS for confirmed cases is $14 billion per year; and if allowance is made for under diagnosed cases, this figure could rise up to $240 billion annually (7–10). The above places IC/BPS among the major health problems that merit research to bridge the wide gaps that exist between diagnosis and treatment. Several treatment regimens have been used to treat this complex condition (11). However, diagnosis is difficult and most treatments show variable, short-term improvements, which are not predictable for individual patients (12) making IC/BPS difficult to treat (13). Thus, there is an unmet need for more effective therapies. Whilst the pathophysiology of IC/BPS is elusive and multi-factorial (14), neurogenic inflammation is believed to important underlying factor contributing to chronic bladder inflammation (14). Histological analysis has shown the urothelium of IC/BPS patients is thin and denuded with or without ulceration (15). This allows harmful urinary constituents to leak into the sub-urothelial layer and depolarize bladder sensory nerve endings (15, 16), resulting in the release of neuroactive substances including substance P (Sub P) and calcitonin gene-related peptide. In addition, the density of SP-positive nerve endings in IC/BPS is not only significantly higher compared to controls but the same nerves are also located in close proximity to mast cells (17).

Sub P activates mast cells resulting in degranulation and the release of both pre-stored and de-novo synthesized pro-inflammatory mediators, which promote chronic bladder wall inflammation (18). These include serine proteases, biogenic amines, arachidonic acid metabolites, cytokines and chemokines (19), which generate chronic inflammation and pain in IC/BPS bladders (20). Upon binding to its G protein-coupled receptors (MRGPRX2), Sub P triggers intracellular signalling events underlying mast cell activation and degranulation.

These include phosphorylation of the enzymes phospholipase C-β (PLC-β), phosphoinositide-3 kinase (PI3K), protein kinase B/Akt, extracellular signal-regulated kinase (ERK1/2) and the signal transducer and activator of transcription 3 (STAT3) (21, 22).

However, the role of neurogenic inflammation in the pathophysiology of the IC/BPS is still contentious. This is due to the uncertainty regarding some factors essential for the development of neurogenic inflammation in the bladders of the patients. Many reports link BPS/IC with a characteristic increase in the density of mast cells in the detrusor layer of the bladders of the patients (23–30). However, other reports stated that there is no significant difference in the mast cell numbers between patients compared to controls (31, 32).

The neurogenic inflammation theory is based on positive responsiveness of mast cells to the neuroactive agents released by the desensitized sensory nerve endings, including the neuropeptide Sub P. Such responsiveness is attributed to mast cell surface expression of the neuropeptide receptor MRGPRX2, which is responsible for Sub P mediated mast cell activation in humans. The expression of MRGPRX2 is variable between different microenvironments (33), and its expression in urinary bladder mast cells has been assessed.

Mānuka honey is derived from the nectar of *Leptospermum scoparium* (Mānuka) trees in New Zealand, and is recognised for a wide range of bioactive properties including its unique non-peroxide antimicrobial activity and biofilm inhibition (34) with recent reports highlighting its potential anti-inflammatory and anti-oxidant effects (35). Interestingly, Medihoney^TM^, a medical-grade Mānuka honey, has been shown to inhibit Ca^++^ ionophore-induced degranulation and histamine release in LAD2 human mast cells (36).

Therefore, the aim of the current study was to further explore the importance of the neurogenic inflammation in the pathophysiology of IC/BPS via assessment of mast cells numbers and their expression of MRGPRX2 in the urinary bladders of BPS/IC patients compared to non-IC/BPS controls using an immunohistochemistry approach. In addition, the second aim was to investigate the potential for Medihoney^TM^ and a sugar-free Mānuka extract to suppress Sub P-induced mast cell degranulation (as a model for neurogenic inflammation), and to explore their potential effects on the activation of MRGPRX2 receptor using representative human cell line models.

## Materials and Methods

### Reagents

100% Medihoney^TM^ (#405, Comvita, New Zealand Ltd), Sub P (# S6883, Sigma), *p*-nitrophenyl-N-acetyl-β-D-glucosaminide (CAS # 3459-18-5, Sigma) and Compound 48/80 (#C2313, Sigma Aldrich).

### Cell culture of the human LAD-2 mast cells

The human mast cell line (LAD2), kindly provided by Professor Arnold Kirshenbaum NIH USA, is a well-established mast cell model system for in vitro neuro-inflammatory testing (37). As per the source instructions, LAD2 cells were cultured in Stem Pro 34-Serum free medium (Invitrogen) enriched with 100 ng /ml recombinant human stem cell factor (Peprotech), 2mM L-glutamine with penicillin/streptomycin (Sigma Aldrich), and 1 ml of Stem Pro growth supplement (Invitrogen) (37). Medihoney^TM^ is kindly provided by Comvita New Zealand Ltd.

### Β-hexosaminidase (degranulation) assay using LAD-2 cells

Following two washes in 1x PBS, the LAD2 cells were re-suspended in Tyrode’s buffer at a density of 50,000 cells per 160 µL of Tyrode’s buffer/well. The cell suspensions were then transferred into a 96-well plate. The LAD2 cells were challenged with 1µM Sub P or 10µg/ml Compound 48/80 (both Compounds being MRGPRX2-specific mast cell activators) for 40-minute with or without 20-minute pre-incubation with Medihoney^TM^ at 1%, 2%, 4% and 6% (W/V) or the Mānuka extract and artificial honey (35% glucose and 35% fructose in ultrapure water), both at 1%, 2%, 4% and 6% (V/V). All agents were prepared in the reaction buffer (Tyrode’s buffer) (38). After incubation, the plates were centrifuged at 693 g for 10 minutes at 4°C. Subsequently, 30µL of each well was transferred into the corresponding well of a 96 well assay plate. A 50 µL aliquot of beta-hexosaminidase substrate (*p*-nitrophenyl-N-acetyl-β-D-glucosaminide (Sigma Aldrich) was added to each well, followed by one hour incubation at 37°C and 5% CO2. Finally, 100 μL of the stop solution 0.2mM glycine was added into each well, and the plate read by a plate reader at Endpoint L1-L2, wavelength 410 to 595 nm.

### The effects of substance P (1uM) and Compound 48/80 (10 ug/ml) on the intracellular Ca++ Flux in MRGPRX2-transfected HEK-293 cells

HEK-293 cells transfected to stably express MRGPRX2 receptor (Creative Biogene) were cultured in high glucose full DMEM medium (Cat# 11965092, ThermoFisher) containing 10% foetal calf serum and 1 μg/mL Puromycin. Cells were cultured to confluence in poly lysine D-coated black/clear bottom 96 well plates. Cells were loaded with Fluo-4 Ca^++^ dye for one hour using Fluo-4 Direct Calcium Assay Kit (Cat# F10471, ThermoFisher), then activated with the MRGPRX2 agonists Sub P (1µm) or Compound 48/80 (10 µg/ml) with or without pre-incubation with Medihoney^TM^, Mānuka extract and artificial honey. The activation-induced rise in the cytoplasmic Ca^++^ levels were assessed using FlexStation II platform (Molecular Devices).

### Immunohistochemistry in IC/BPS urinary bladder biopsies

The study commenced after obtaining ethical approval of the Research Ethics Committee (REC) (IRAS ID 246784). Formalin-fixed paraffin embedded (FFPE) urinary bladder biopsies from 17 IC/BPS patients and 16 non-IC/BPS controls were involved into the study. The included FFPE biopsies were taken for diagnostic purposes after patient consent. IC/BPS diagnosis was based according to the 2008 ESSIC diagnostic criteria. The control group include healthy bladder tissue from carcinoma in situ patients or patients with non-specific bladder wall inflammation. 4 serial sections, each of 4μm thickness, were cut from each bladder biopsy and stained with antibodies against mast cell-specific proteases tryptase (AA1) and chymase (CC1), as unique mast cell markers, as well as anti-MRGPRX2 (Abcam, USA). All staining was performed using an automated auto Stainer platform. Following microscopic scanning, images from the serial sections were aligned and superimposed using Adobe Photoshop CS6 software. Mast cell quantification and co-expression patterns were studied using Image J software. Statistical differences between the IC/BPS and the control groups were analysed by Mann Whitney U test and unpaired t-tests using GraphPad Prism 9 software.

### Statistical Analysis

Data were included in a database and analysed using GraphPad Prism 9 software (GraphPad Software, San Diego, USA). Data normality was tested using the Shapiro-Wilk method. Results which were normally distributed were expressed as mean ± SEM, where differences between two or more groups were assessed by one-way ANOVA. Results which were not normally distributed are expressed as median and interquartile range, where differences between two or more groups were assessed by Kruskal-Wallis and Mann-Whitney U tests. A *p value of <0.05 2-sided* was taken as statistically significant.

## Results

### The effect of Medihoney^TM^ on LAD-2 cells

After one hour incubation of the LAD2 cells with 2%, 4% or 6% concentrations of Medihoney^TM^, β-hexosaminidase levels remained similar to the spontaneous levels, suggesting that Medihoney^TM^ alone at all concentrations used did not induce LAD2 cell degranulation. On the other hand, the MRGPRX2 agonists, Sub P and Compound 48/80, induced 42% and 61% release, respectively. The non-specific mast cell activator Ca++ ionophore was included as a positive control and induced 86% release above spontaneous. The overall cellular content of β-hexosaminidase (total) was prepared by completely lysing the cells in 1% Triton-X in 1X PBS (Figure 1 (A)).

**Figure 1:**
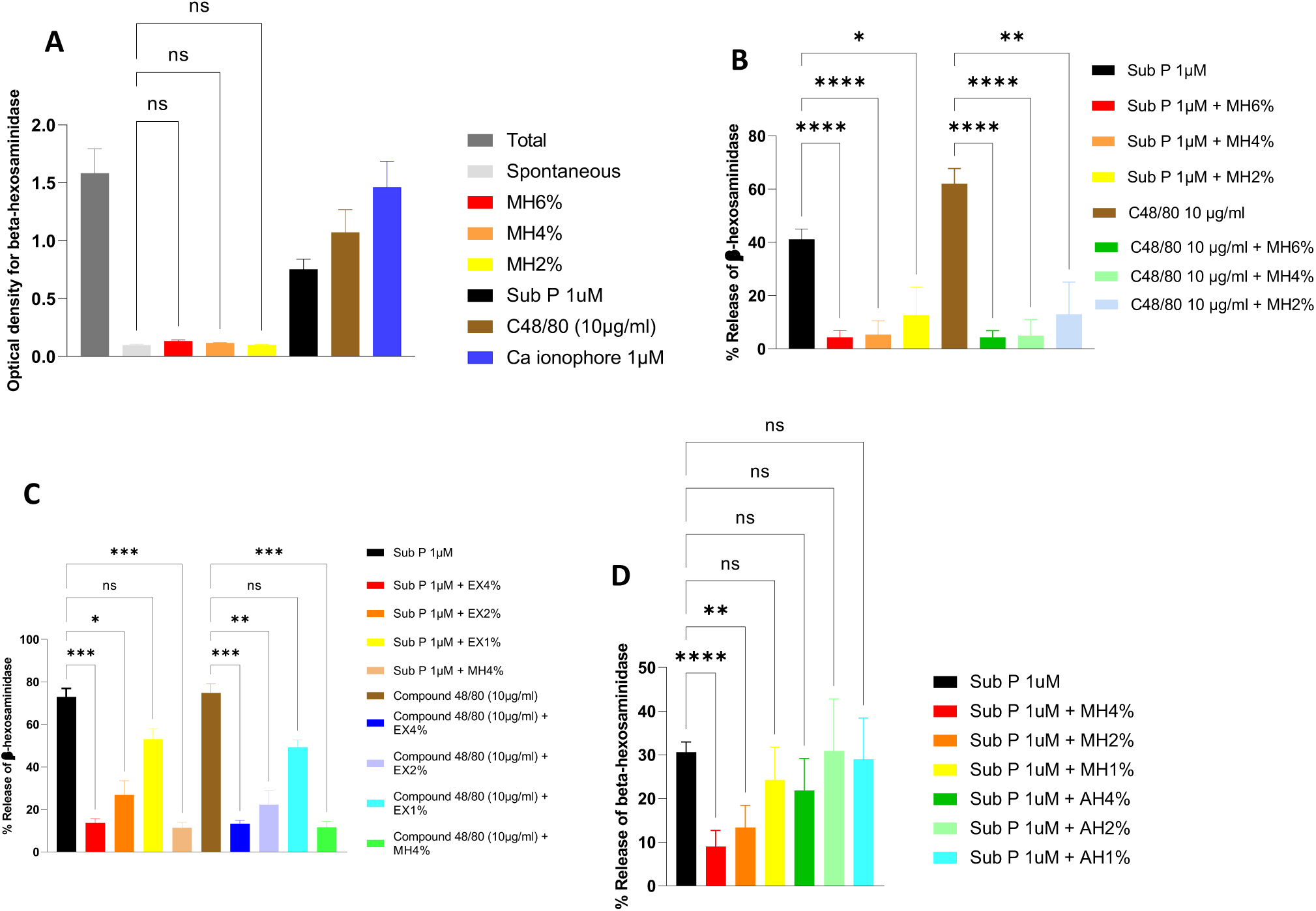
The effect of Medihoney^TM^ on β-hexosaminidase from LAD2 cells. (A) Optical density for the release of β-hexosaminidase by LAD2 cells following one-hour incubation with Medihoney^TM^ (MH) at 2, 4% and 6 %, 1μM substance P (Sub P), 10μg/ml Compound 48/80 (C48/80) and 1µM Ca^++^ ionophore (Positive control). (B) & (C) represent the percentage release of β-hexosaminidase induced by 40-minute incubation with 1μM Sub P and 10 μg/ml Compound 48/80, respectively, with or without pre-incubation with Medihoney^TM^ (MH) or the Mānuka extract (EX) at 2%, 4% and 6 % in LAD2 cells, while (D) represents the Sub P (1μM)-induced percentage release with or without pre-incubation with artificial honey (MH) at 1%, 2% and 4 %. The mean ± SEM for the spontaneous release was 6.7 % ± 0.09 % of the total value. The overall cellular content of β-hexosaminidase (total) was prepared by completely lysing the cells in 1% Triton-X in 1X PBS. Each bar represents mean ± SEM (** P value ˂0.01, *** P value ˂0.001 and **** P value ˂0.0001, n=12). The graph represents the results of 4 independent assays, and each assay was performed in triplicates.

### The effect of Medihoney^TM^ on substance P and Compound 48/80-induced LAD2 cell degranulation

Pre-incubation with Medihoney^TM^ at 2%, 4% and 6% resulted in 79%, 97% and 94% inhibition of the β-hexosaminidase release induced by both Sub P or Compound 48/80, respectively (Figure 1 (B) & (C)). Therefore, 2% and 4% where chosen for subsequent experiments. Importantly, pre-incubation with artificial honey at 2% and 4% did not inhibit the β-hexosaminidase release induced by Sub P (Figure 1 (D)). This finding excludes the sugar content from being responsible for the mast cell stabilizing effects of Medihoney^TM^.

### The effects of substance P (1µM) and Compound 48/80 (10 µg/ml) on the intracellular Ca++ Flux in MRGPRX2-transfected HEK-293 cells

Sub P (1µM) and Compound 48/80 (10 µg/ml) induced a significant increase in the intracellular Ca^++^ levels in the MRGPRX2-transfected HEK-293 cells within 18 seconds of treatment. However, both failed to induce any Ca^++^ signalling in the standard non-transfected HEK-293 cells. Typically, as a non-specific Ca^++^ ion carrier, Ca^++^ ionophore 1µM induced a rise in the intracellular Ca^++^ levels in both the transfected and non-transfected cells. The Sub P vector (0.1M acetic acid) failed to induce any increase in the intracellular Ca^++^ levels in HEK-293 cells or degranulation in LAD2 cells (Figure 2).

**Figure 2:**
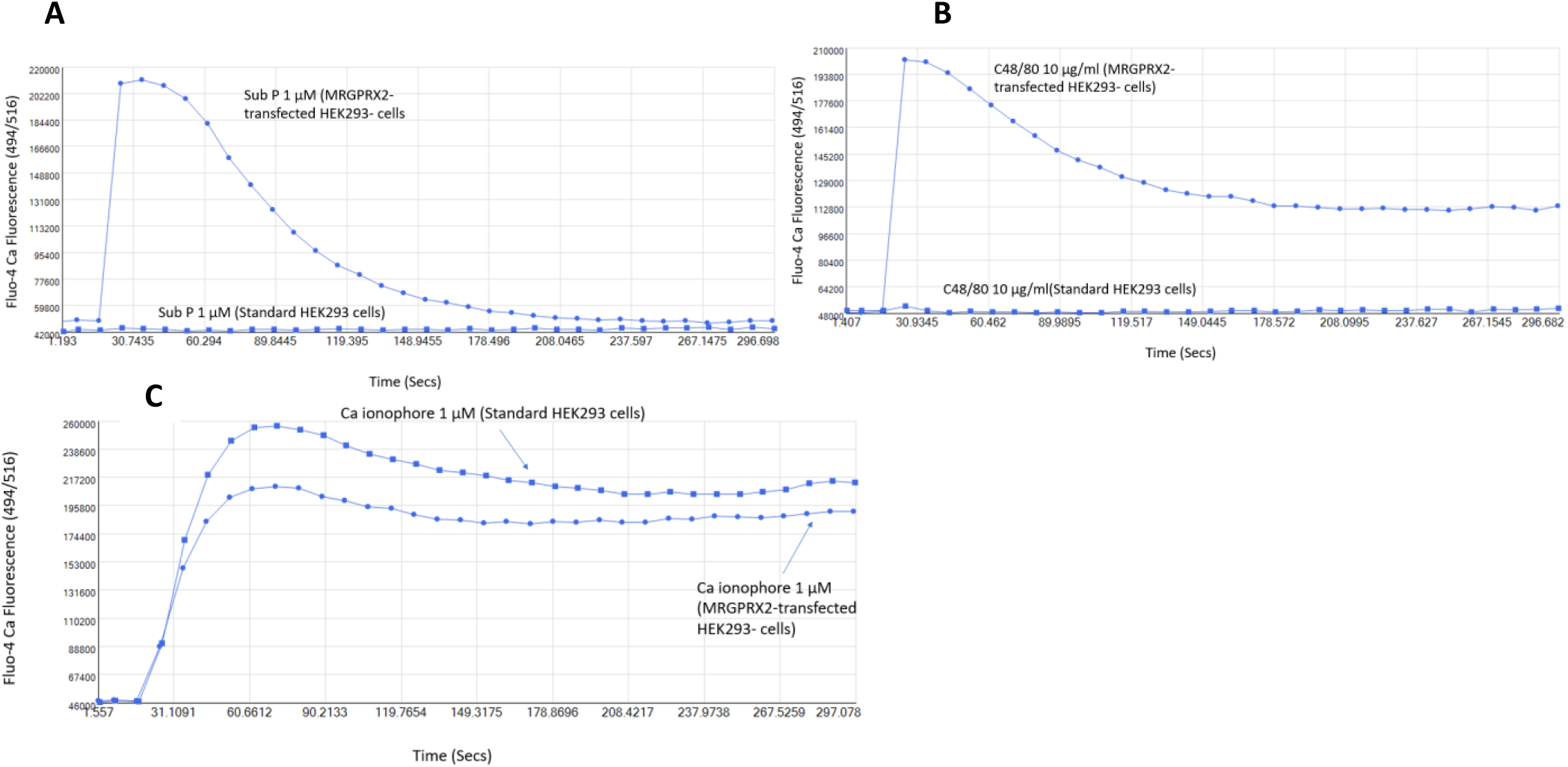
The effect of substance P and Compound 48/80 on the intracellular Calcium signalling in MRGPRX2-transfected HEK-293 cells. Intracellular Calcium levels measured by Fluo-4 imaging in MRGPRX2-transfected HEK-293 cells (A) vs standard non-transfected HEK-293 cells (B) upon challenge with 1µM Sub P (A), 10µg/ml Compound 48/80 (B) and 1µM Ca ionophore (C).

### The effect of Medihoney^TM^ on the substance P-induced rise in the cytoplasmic Ca^++^ levels in MRGPRX2-expressing HEK-293 cells

Twenty-minute pre-incubation of the HEK-293 cells with Medihoney^TM^ or the Manuka honey extract at 2 and 4% induced dose-dependent inhibition of the Sub P (Figure 3 (A, E, H & J)) or the Compound 48/80-induced Ca^++^ flux (Figure 3 (B, D, G, & I)), while artificial honey (AH) at 2% and 4% did not inhibit the Sub P-induced Ca^++^ flux (Figure 3 (C &H)).

**Figure 3:**
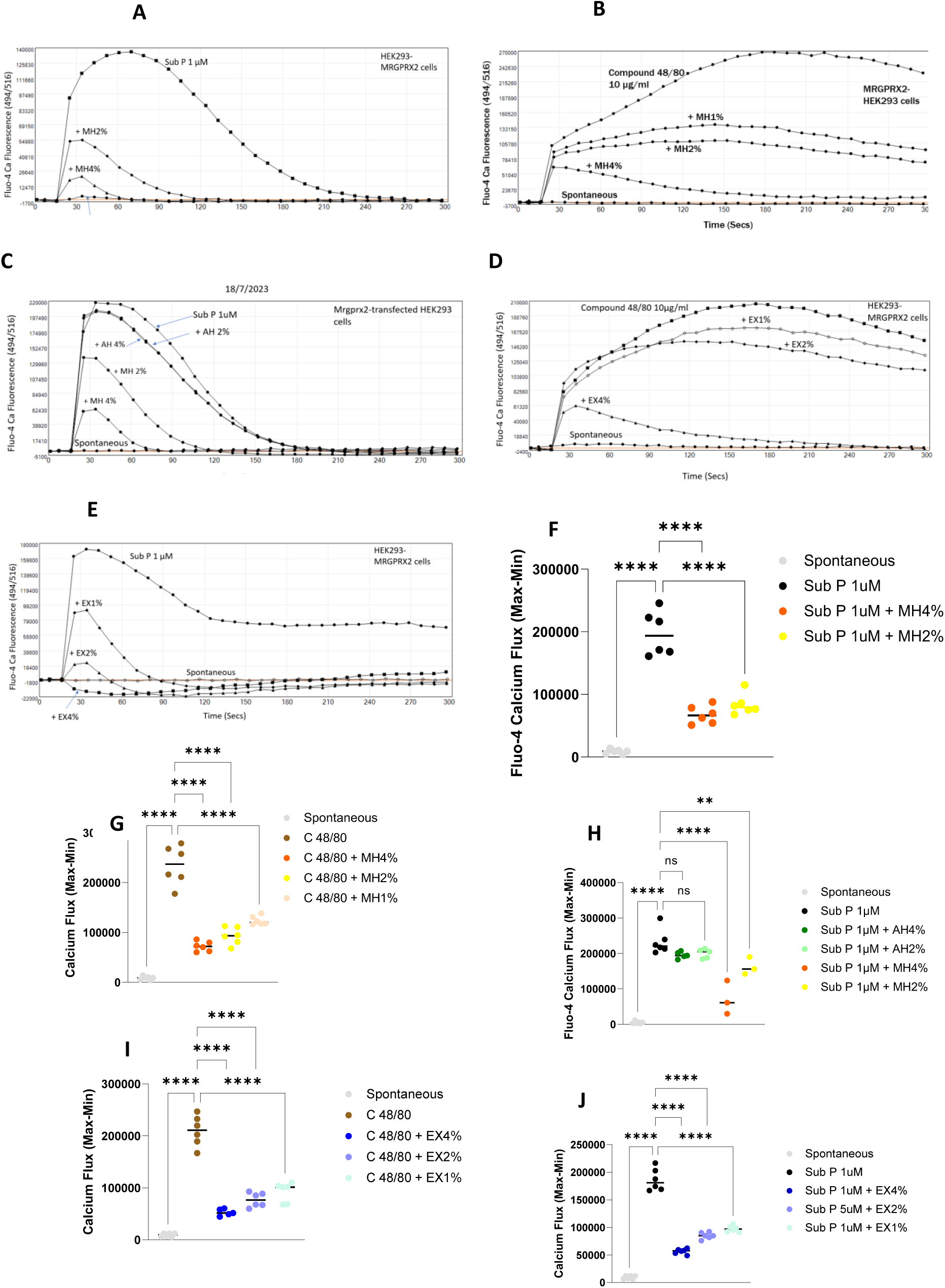
The effect of Medihoney^TM^ on the substance P and Compound 48/80-induced intracellular Calcium signalling in MRGPRX2-transfected HEK-293 cells. Intracellular Calcium levels measured by Fluo-4 imaging in MRGPRX2-transfected HEK-293 cells upon challenge with 1µM Sub P with or without pre-incubation with Medihoney^TM^ (MH) 2% and 4% (A &F), artificial honey (AH) 2% and 4% (C & H), or Mānuka extract (EX) 2% and 4% (E & J). (B & G) and (D & I) represent the intracellular Calcium levels measured by Fluo-4 imaging in MRGPRX2-transfected HEK-293 cells upon challenge with 10 µg/ml Compound 48/80 or without pre-incubation with Medihoney^TM^ (MH) 2% and 4% and Mānuka extract (EX), respectively, (** P˂0.01 and **** P˂0.0001 (n=8)).

**Figure 4:**
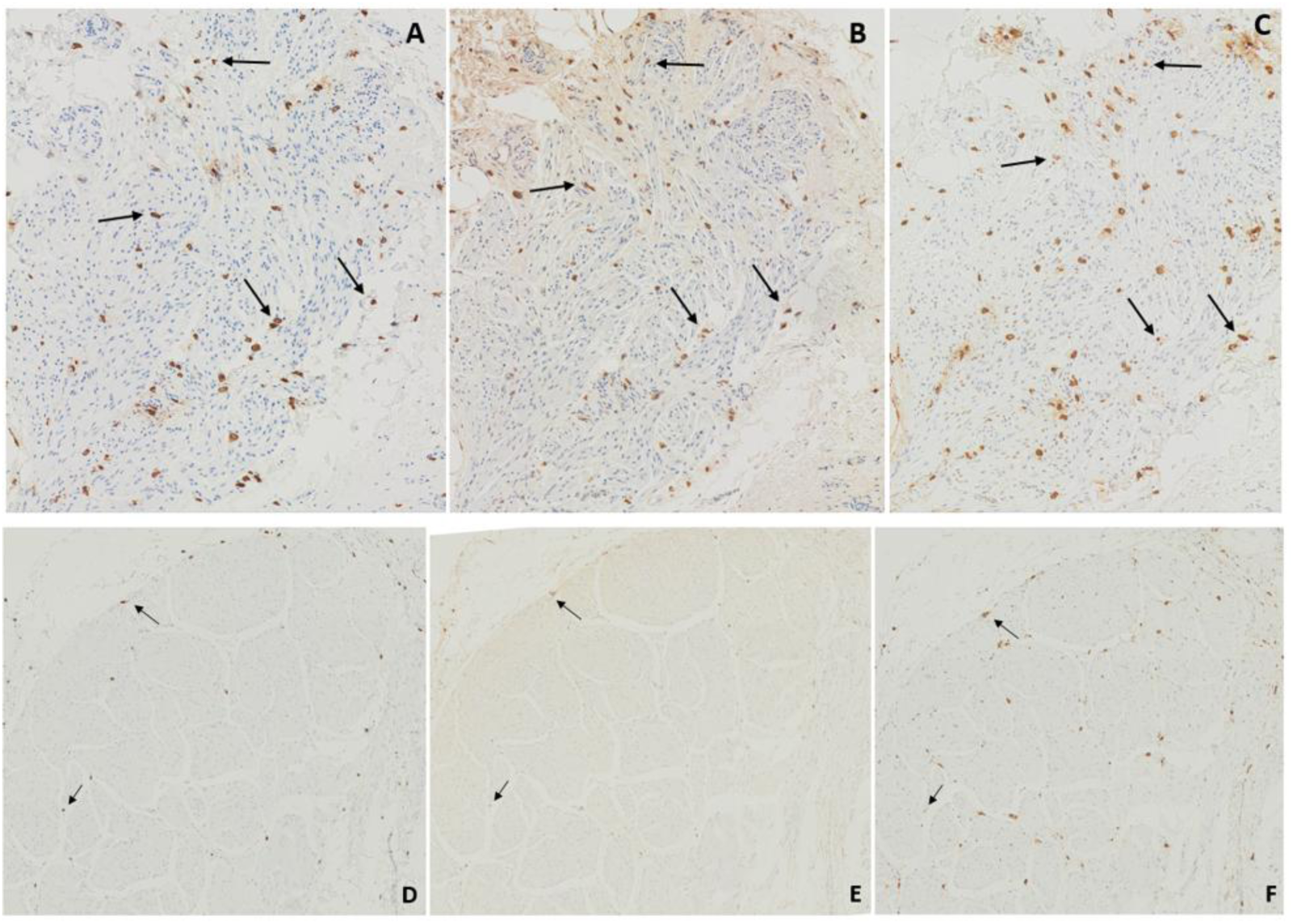
Immunohistochemical analysis of the MRGPRX2 expression by the mast cells in the detrusor layer urinary bladders of IC/BPS vs controls. Serial sections of 2 urinary bladder biopsy from an BPS/IC patient (A, B and C), and healthy bladder tissue control (D, E and F) showing mast cells in the detrusor layer, some are indicated in arrows, stained with antibodies for mast cell chymase (A & D), MRGPRX2 (B & E) and mast cell tryptase (C& F).

**Figure 5:**
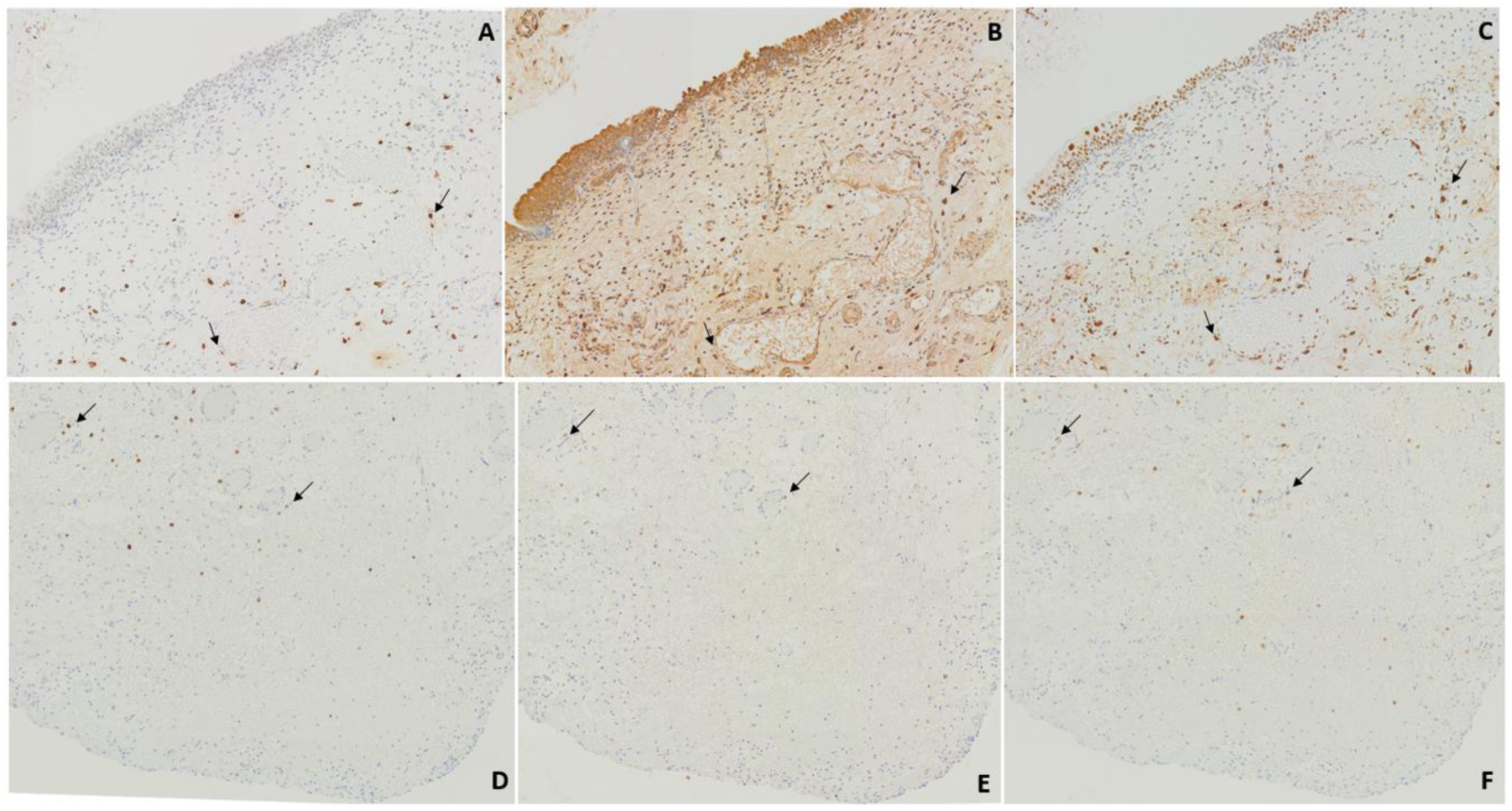
Immunohistochemical analysis of the MRGPRX2 expression by the mast cells in the lamina propria urinary bladders of IC/BPS vs controls. Serial sections of 2 urinary bladder biopsy from an BPS/IC patient (A, B and C), and healthy bladder tissue control (D, E and F) showing mast cells in the lamina propria, some are indicated in arrows, stained with antibodies for mast cell chymase (A & D), MRGPRX2 (B & E) and mast cell tryptase (C& F).

### Immunohistochemical analysis of mast cell density and MRGPRX2 expression in FFPE bladder biopsies from IC/BPS patients compared to controls

#### Mast cell density and subtypes

In the BPS/IC group, mast cells were abundant in both detrusor layer and the lamina propria (LP). The mean (± SEM) density of mast cells stained with tryptase (all mast cells) was 166.1 ± 21.02 and 215.0 ± 19.05 cells/mm^2^ in the detrusor layer and the lamina propria, respectively. This was significantly higher compared to the control group (72.64 ± 9.150 and 84.88 ± 9.212, respectively; Figure 6 (A&B)). Similarly, the mean (± SEM) density of the Chymase^+ve^ mast cells (MC_TC_) in the IC/BPS group was 155.7 ± 23.86 and 181.6 ± 17.09 cells/mm^2^, in the detrusor layer and the LP, respectively. This was significantly higher compared to the control group (59.83 ± 9.153 and 65.27 ± 8.710, respectively; Figure 6 (C&D)). In the BPS/IC group, the mean (± SEM) percentage of chymase expression by the Tryptase^+ve^ mast cells was 70.95 ± 5.371 and 84.47 ± 4.209, in the detrusor and LP, respectively. This was significantly higher compared to the control group (54.45 ± 4.546 and 55.21 ± 4.332, respectively; Figure 72), indicating an IC/BPS-related class switch towards the MC_TC_ subtype.

**Figure 6:**
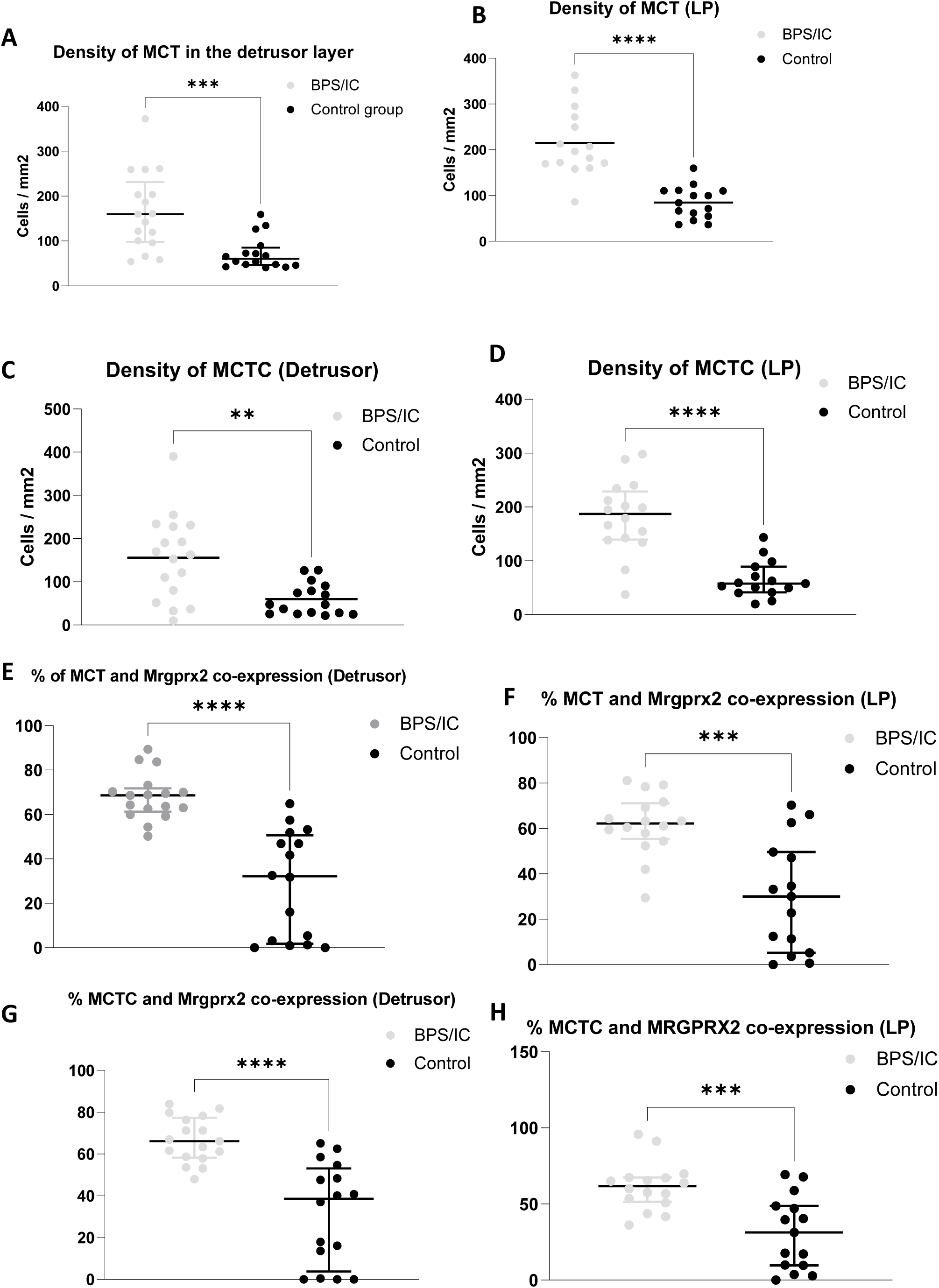
Graphical representation of the mast cell densities and their MRGPRX2 expression in the urinary bladders of IC/BPS patients vs controls. (A) & (B) are scatter plots representing the density of mast cells stained with tryptase (MCT) in the urinary bladders of the IC/BPS patients vs controls in the detrusor vs the LP layer, respectively. (C) & (D) are scatter plots representing the density of mast cells stained with Chymase (MCTC) in the urinary bladders of the IC/BPS patients vs controls in the detrusor vs the LP layer, respectively. (E) & (F) Scatter plot representing the percentage of MRGPRX2 expression by the Tryptase^+ve^ mast cells (MCT) in the urinary bladders of IC/BPS patients vs controls in the detrusor and LP layers, respectively. (G) & (H) Scatter plot representing the percentage of MRGPRX2 expression by the Chymase^+ve^ mast cells (MCTC) in the urinary bladders of IC/BPS patients vs controls in the detrusor and LP layers, respectively. Each bar represents the mean, (**P˂0.005, *** P value 0.002 & **** P value ˂0.0001).

#### Mast cell expression of MRGPRX2 in the patients vs controls

In the BPS/IC group, the mean (± SEM) percentage of tryptase+ve mast cells expressing MRGPRX2 was 67.99 ± 2.537 and 61.75 ± 3.366 in the detrusor and LP layers, respectively. This was significantly higher compared to the control group (28.38 ± 6.004 and 29.97 ± 6.352, n the detrusor and LP layers, respectively; Figure 6E-F)). Similarly, in the BPS/IC patient group, the mean (± SEM) percentage of Chymase^+ve^ mast cells expressing MRGPRX2 was 66.66 ± 2.629 and 61.64 ± 3.975 in the detrusor and LP, respectively. This was significantly higher compared to the control group (31.44 ± 6.046 and 24.99 ± 7.296, in the detrusor and LP, respectively; Figure 6 (G&H). This indicates an IC/BPS-related increase in MRGPRX2 expression by urinary bladder mast cells.

## Discussion

Mast cells are key effectors in both the innate and adaptive immune responses. Their abnormal activation is believed to be involved in both the allergic and pseudo-allergic reactions implicated in many chronic inflammatory conditions (19). A mutual activation cycle between the neuropeptide Sub P releasing nerve endings and tissue resident mast cells is believed to maintain the chronic inflammatory state in interstitial IC/BPS. Such interaction has been implicated in the pathophysiology of many other chronic inflammatory diseases including atopic dermatitis, chronic urticaria, rheumatoid arthritis, and severe asthma (39). Mast cell responsiveness to the neuropeptide Sub P in humans depends on their expression of MRGPRX2, a recently identified G protein coupled receptor (40). The expression of such a receptor on mast cells varies between different tissue micro-environments and its expression on urinary bladder mast cells has not been explored previously (33).

In the current study, the average density of mast cells stained with tryptase or chymase in both the detrusor or the LP layer of the BPS/IC group was significantly higher compared to the control tissue. Such evidence is consistent with previous reports highlighting the increased numbers of the tissue resident mast cells in the bladders of BPS/IC patients (29, 41–43). Regarding mast cell subtypes, about 70% and 84% of the Tryptase^+ve^ mast cells in the detrusor and the LP layer of the BPS/IC group co-expressed chymase, which was significantly higher than controls (only 50% co-expression). Such evidence suggests a shift of the mast cell subtype from MC_T_ to MC_TC_ in the BPS/IC group, which is consistent with other reports highlighting the shift of the MC subtype from MC_T_ to MC_TC_ in the pulmonary mast cells of severe asthma patients (44). Based on previous reports highlighting the pro-inflammatory and cell lysis effects of mast cell chymase (45, 46), the shift in the mast cell subtype to MC_TC_ might, in part, explain the presence of the urothelial barrier defect seen in the bladders of the BPS/IC patients.

MRGPRX2 was expressed by around 60% and 70% of the mast cells in the LP and the detrusor layers of the IC/BPS bladders, which was significantly higher compared to controls (30% co-expression). To our knowledge, this is the first evidence confirming the expression of MRGPRX2 on the mast cells in a urinary bladder tissue and, specifically, in BPS/IC patients. Nevertheless, such evidence indicates the upregulation of MRGPRX2 receptor expression by mast cells in BPS/IC. In the current study, MC_TC_ was the dominant subtype (around 70% of all mast cells). This finding is supported by the MRGPRX2 expression, which is believed by many to be exclusively expressed by connective tissue mast cells (MC_TC_) (44, 47).

Medihoney^TM^ has been shown to inhibit mast cell degranulation and histamine release induced by Ca++ ionophore, a non-specific strong inducer of mast cell activation in vitro (20). In the current study, both Medihoney^TM^ and a sugar-free Mānuka honey extract inhibited mast cell degranulation (as assessed by β-hexosaminidase release) induced by the neuropeptide Sub P in a dose-dependent manner, suggesting a strong potential for these Mānuka-derived agents to supress neurogenic mast cell activation and tissue inflammation. Maximal inhibition (97%) was achieved using 4% of Medihoney^TM^ or the Mānuka Extract (Figure 1).

Such an effect was accompanied by dose-dependent inhibition of the Sub P-induced Ca^++^ signalling in MRGPRX2-transfected HEK-293 cells, indicating inhibition of the receptor activation. Such evidence strongly supports an anti-inflammatory role of these Mānuka- derived agents against neurogenic inflammation. The artificial honey with comparative sugar levels to Mānuka honey did not inhibit mast cell degranulation or the MRGPRX2-mediated Calcium internalisation. This suggests that the anti-inflammatory effect of Medihoney^TM^ is independent from its sugar content. Such evidence is further supported by the fact that the sugar-free Mānuka honey extract replicated the same effects of Medihoney^TM^ on both degranulation and Calcium internalisation (Figures 1&3). Thus, the Mānuka extract could potentially be suitable alternative to Medihoney^TM^ for use in diabetic patients with chronic inflammatory conditions e.g., IC/BPS, atopic dermatitis, etc.

Unlike other anti-inflammatory agents, honey is not a pure agent, it has a unique natural composition consisting of a mixture of biologically active agents. This enables honey to produce several biological effects. For example, it possesses strong antimicrobial activity against a broad spectrum of bacterial strains including those that are resistant to antibiotics (48), and strongly inhibits bacterial biofilm formation (34). Furthermore, it promotes angiogenesis and tissue regeneration (49) and inhibits inflammation independent of its anti-microbial properties. In addition, Medihoney^TM^ was well tolerated by urothelial cells and protected them from cytotoxic insults while maintaining urothelial barrier integrity in IC/BPS bladder models (50). Taken together, the above studies demonstrate that Medihoney^TM^ possesses three properties that could be exploited clinically. Thus, it is anti-microbial, pro-angiogenic and anti-inflammatory. This “triple action” would be ideally suited for the treatment and management of IC/BPS.

### Conclusion and applications

Our current findings demonstrate the potential anti-inflammatory properties of Medihoney^TM^, through its ability to inhibit mast cell degranulation induced by the neuropeptide Sub P. Moreover, the study provides the first evidence of the modulatory effects of Medihoney^TM^ on the intracellular signalling events accompanying mast cell activation.

These data support the potential use of Medihoney^TM^ as an anti-inflammatory agent in neurogenic inflammatory conditions such as IC/BPS and atopic dermatitis. Specifically, aqueous solutions of Medihoney^TM^ could be used intravesically to aid the management of patients with IC/BPS.

This work was partially supported by Comvita through the provision of Medical Grade Mānuka Honey (Medihoney™), a sugar-free Mānuka honey extract, and product-related information. Comvita also contributed towards funding a postdoctoral research fellowship. The company had no involvement in the design, conduct, analysis, interpretation, or reporting of this study.

## Acknowledgements

The authors gratefully acknowledge Comvita for providing Medical Grade Mānuka Honey (Medihoney™), the sugar-free Mānuka honey extract, and product-related information that supported this research. The authors also acknowledge Comvita for its contribution towards funding a postdoctoral research fellowship.

